# Ontogeny and Trophic Factor Sensitivity of Gastrointestinal Projecting Vagal Sensory Cell Types

**DOI:** 10.1101/2022.07.04.498754

**Authors:** Meaghan E. McCoy, Anna K. Kamitakahara

## Abstract

Vagal sensory neurons located in the nodose ganglion provide a direct line of communication from the gut to the brain. Information, such as stomach stretch or the presence of ingested nutrients in the intestine, is conveyed to the caudal medulla via specialized cell types that express unique marker genes. Here, we leverage vagal cell type marker genes identified in adult mice to determine when specialized vagal subtypes arise developmentally, as well as the factors that shape their growth and target innervation. Initial sequencing and pathway analysis comparing early postnatal to adult nodose ganglia identified an enrichment of differentially expressed genes (DEGs) related to axon growth and guidance, synaptic strengthening, and DEGs downstream of brain derived neurotrophic factor (BDNF). Subsequent experiments performed to screen for trophic factor sensitivity revealed that both BDNF and glial cell derived neurotrophic factor (GDNF) robustly stimulate neurite outgrowth from early postnatal nodose ganglion explants *in vitro*. Anatomical examination of whole-body trophic factor expression in perinatal mice revealed that BDNF is expressed by neurons of the nodose ganglion itself, while GDNF is expressed by developing intestinal smooth muscle cells. Thus, BDNF may support vagal neurons as a locally derived trophic factor, while GDNF may act as a target derived trophic factor supporting the growth of vagal sensory processes at distal innervation sites in the gut. Consistent with this, expression of the GDNF receptor, *Gfra1*, was enriched in vagal afferent cell types that project to the gastrointestinal tract. The BDNF receptor, *Ntrk2,* was expressed by the majority vagal sensory neurons, irrespective of cell type. Lastly, spatial mapping of genetic markers in the nodose ganglion demonstrates that defined vagal cell types begin to emerge as early as embryonic day 13, even as sensory neurons continue to grow to reach gastrointestinal targets. Despite the early onset of expression for some cell type marker genes, expression patterns of many individual cell type markers appear largely immature in prenatal life and mature considerably by the end of the first postnatal week. Together, the data support a role for GDNF in stimulating vagal sensory growth to gastrointestinal targets and establish a prolonged perinatal timeline for vagal sensory cell type maturation.

## Introduction

Abdominal-projecting vagal sensory neurons located in the nodose ganglion convey vital information from the gut to the brain used to regulate feeding behavior and coordinate descending motor control to the pancreas and gastrointestinal (GI) tract (Berthoud and Neuhuber, 2000; Schwartz, 2000; Travagli and Anselmi, 2016). GI-related sensory information is transmitted via specialized cell types that detect stomach or intestinal stretch, the presence of ingested nutrients, and/or the presence of microbial pathogens and inflammatory signals (Phillips and Powley, 1996; Wang and Powley, 2000; Powley and Phillips, 2004; Williams et al., 2016a; Bellono et al., 2017; Beutler et al., 2017; Han et al., 2018; Kaelberer et al., 2018; Bai et al., 2019; Kupari et al., 2019a; Ye et al., 2020). The sensory modalities and properties of individual vagal afferent cell types have been defined anatomically, morphologically, electrophysiologically, functionally, and at the transcriptome level in rodents. However, the vast majority of studies have focused on the properties of adult vagal neurons, providing only a limited understanding of how and when specific vagal cell types arise and mature during embryonic and early postnatal life, and the factors that guide these developmental processes.

Published studies that have examined vagal development provide a basic framework for understanding the trajectory of neurodevelopmental events from neurogenesis to functional circuits. Vagal afferent neurons are derived from the third epibranchial placodes and join together with neural crest-derived jugular ganglion neurons to form a fused jugular-nodose complex (Douarin et al., 1986; Baker and Bronner-Fraser, 1997; Vermeiren et al., 2020). Only the ventrally positioned neurons of the nodose ganglion project to the GI tract. As early as embryonic day (E) 10.5 in mice, vagal sensory neurons begin to extend projections along the esophagus (Ratcliffe et al., 2006, 2011; Niu et al., 2020). Projections then grow rapidly to reach the stomach from E11.5 to E13.5 and reach the intestines between E14.5 and 16.5 (Ratcliffe et al., 2006, 2011; Niu et al., 2020). Intestinal morphology develops in concert with innervation. At E14.5 intestinal villi have yet to form, with few neuronal projections observed to enter the stromal space (Hao et al., 2020). Villi begin to emerge from E15.5 to E16.5, during which time neural projections enter the stroma to span the entire length of the villus (Hao et al., 2020).

Despite extending the entire length of the GI tract prenatally, vagal projections mature considerably during the first few postnatal weeks (Smith, 2006; Murphy and Fox, 2007). For example, this has been shown in mechanosensitive vagal sensory axons specialized to detect GI stretch that form intraganglionic laminar endings (IGLEs) and intramuscular arrays (IMAs). The innervation patterns of these neurons continue to mature morphologically throughout the first postnatal week in rodents, approaching the density of terminal structures observed in the adult GI tract by postnatal day (P) 8 (Murphy and Fox, 2007). At this time, gastric distention clearly inhibits feeding behavior in early postnatal rats (Phifer et al., 1986). Interestingly, the ability of ingested nutrients to inhibit feeding behavior does not arise until slightly later (Phifer et al., 1986; Weller, 2000). This is consistent with data demonstrating separate subpopulations of extrinsic vagal and spinal neurons that are mechanosensitive (e.g. vagal IGLEs and IMAs) or nutrient sensitive (e.g. vagal and spinal mucosal projecting) (Phillips and Powley, 1996; Powley, 2000; Wang and Powley, 2000; Williams et al., 2016a; Beutler et al., 2017; Bai et al., 2019; Kaelberer et al., 2020; Goldstein et al., 2021), although, polymodal mechano- and chemo-sensitive cell types also likely exist (Bai et al., 2019; Kupari et al., 2019b). Studies using gastric preloads to limit or avoid the effects of oral gustatory and somatosensory stimulation and taste have further established a timeline for the development of postingestive inhibition of feeding in response to nutrients, such as carbohydrates, fats, and proteins. In rats, gastric preloads of glucose, corn oil, or peptone do not appreciably inhibit independent food intake compared to non-nutritive preloads until after the second postnatal week (~P15-18) (Weller, 2000; Weller et al., 2000; Smith, 2006; Davis et al., 2013). Together, the anatomical and behavioral data provide essential temporal information on the developmental trajectory of vagal sensory development.

The mechanisms guiding these critical developmental events remain only partially defined. Early neuronal survival and growth of subpopulations of vagal neurons are known to depend upon the support of brain derived neurotrophic factor (BDNF) and glial derived neurotrophic factor (GDNF). Congenital deletion of BDNF results in a 65% reduction in the number of neurons in the nodose ganglia by P0 (Ernfors et al., 1994; Hellard et al., 2004). Similarly, deletion of GDNF results in a 40% reduction in the number of nodose ganglion neurons by P0 (Moore et al., 1996). It remains to be determined which subpopulations of vagal cell types are affected by each trophic factor developmentally. Both BDNF and GDNF have been reported to stimulate neurite outgrowth in cultured vagal sensory neurons (Niwa et al., 2002). Furthermore, the density of IGLEs in the stomach are reduced following BDNF deletion, providing evidence that BDNF may support vagal cell types that project to the GI tract (Murphy and Fox, 2010). It is unknown whether other vagal subpopulations are similarly responsive, and experiments to address this are complicated by the fact that BDNF deletion is lethal. Recent single cell sequencing experiments have suggested that up to 27 unique vagal sensory cell types can be defined in adult mice (Bai et al., 2019; Kupari et al., 2019b). These molecularly-defined cell type markers can be used to further identify which vagal subpopulations are affected by specific trophic factors as well as the developmental time points at which cell type specific populations arise.

The present study used bulk RNA sequencing and pathway analysis combined with *in vitro* explant studies and *in situ* hybridization to further illuminate the molecular pathways and processes at play as vagal afferent neurons develop. Our novel results reveal important relationships between trophic factors and specific vagal subpopulations and provide critical information on the timing of vagal cell type maturation.

## Methods

### Animals

Animal care and experimental procedures were performed in accordance with the Institutional Animal Care and Use Committee of The Saban Research Institute, Children’s Hospital Los Angeles. Mice were housed in the vivarium on a 13:11 hour light:dark cycle (lights on at 06:00 hours, lights off at 19:00 hours) at 22°C with ad libitum access to a standard chow diet (PicoLab Rodent Diet 20, #5053, St. Louis, MO). The C57BL/6J, *Phox2b^cre^*, and cre-dependent TdTomato reporter lines (*TdTomato*) were obtained from The Jackson Laboratory (C57BL/6J strain 000664, RRID:IMSR_JAX:000664; B6(Cg)-Tg(Phox2b-cre)3Jke/J stain 016223, RRID:IMSR_JAX:016223; and B6.Cg-Gt(ROSA)26Sortm14(CAG-tdTomato)Hze/J, stock 007914; Ai14, RRID:IMSR_JAX:007914, respectively). All lines were bred and maintained in the Saban Research Institute vivarium on a C57BL/6J background.

### RNA Sequencing

Tissue from the nodose ganglion was dissected from male mouse pups on postnatal day (P) 7 and from male and female 12-week-old C57BL/6J mice. While the nodose ganglion is fused together with the jugular ganglion, transection of the fused ganglia at the narrowing between the two facilitated enrichment of the sample by neurons of nodosal origin. Each sample consisted of the left and right nodose ganglia from a single mouse obtained from independent litters.

RNA was isolated from each sample using the QIAGEN RNeasy micro kit as specified by the manufacturer. Total RNA quality was determined by measuring RNA Integrity Number (RIN) on an Agilent Bioanalyzer RNA Pico chip. Using 50 micrograms of RNA, transcriptome libraries were prepared and processed with NEB Next Ultra II Directional RNA Library Prep Kit with single indexing following manufacturer’s protocol. Library quality was determined using an Agilent Bioanalyzer DNA 1000 chip. Libraries were sequenced using paired end (150 bp × 150 bp) chemistry on the Illumina HiSeq platform (350 M reads).

Reads were mapped to the UCSC mm10 transcript set using Bowtie2 version 2.1.0 and gene count was estimated using RSEM v1.2.15. The TMM method was used for gene count normalization and differentially expressed genes were identified using the edgeR R program. Genes exhibiting fold-changes greater than 1.5 with false discovery rate adjusted p-values < 0.05 were considered differentially expressed.

Raw and processed data have been deposited in the Gene Expression Omnibus (GEO) and are available under accession number GEO:XXXXX

### Ingenuity Pathway Analysis

Differentially expressed genes (DEGs) between P7 and 12-week male samples were further analyzed using QIAGEN’s Ingenuity Pathway Analysis (IPA) software (Fall 2020 release). To restrict the analysis to the highest confidence DEGs, cutoffs for average expression counts greater than 5, expression z-scores less than 2, fold-changes greater than 1.5, and adjusted p-values below 0.05 were used. Using this list of analysis ready molecules, statistically significant canonical pathways and upstream regulators were identified.

### Immunohistochemistry

Tissue for immunofluorescence staining was collected on embryonic days (E) 13 and 15, and from P0 and 7 mice. For embryonic tissue collection, timed pregnant breeding pairs were set, with the day of vaginal plug detection designated as E0. Sex was not determined for embryonic samples. Embryonic and P0 tissue was collected and immersed overnight in fixative (4% paraformaldehyde in 0.1M phosphate-buffered saline (PBS, pH 7.4)). P7 mice were deeply anesthetized by intraperitoneal injection of ketamine:xylazine (100mg/kg:10mg/kg, Henry Schein, Melville, NY) and perfused transcardially with 0.9% saline, followed by fixative. Collected tissues were postfixed for two hours, cryoprotected overnight in 20% sucrose in PBS, embedded in Tissue-Tek® Optimal Cutting Temperature Compound, and frozen over liquid nitrogen vapors or powdered dry ice. From whole embryos or dissected nodose ganglion tissue, 20μm-thick cryostat sections were collected in five series representing the entire width of the nodose ganglion. For whole embryos, sections were collected in the sagittal plane. For dissected nodose ganglion tissue, sections were collected longitudinally. Slides were stored at −20°C until processed.

For immunofluorescence labeling, slides were incubated for one hour in blocking buffer containing 5% normal donkey serum (Jackson ImmunoResearch, West Grove, PA) and 0.3% Triton X-100 in PBS, then overnight in antibody solution containing 2% normal donkey serum and 0.3% Triton X-100 in PBS with primary rabbit anti-activated caspase 3 antibody (Cell Signaling, Cat# 9661). Sections were washed five times for five minutes in PBS, then incubated in secondary Alexa Fluor^®^ 594 AffiniPure F(ab’)_2_ Fragment Donkey Anti-Rabbit IgG (Jackson ImmunoResearch Cat# 711-586-152, RRID:AB_2340622). After several rinses in PBS, sections were counterstained with DAPI (Thermo Fisher Scientific Cat# D1306), and a coverslip was applied using ProLong Gold Antifade Mountant (Thermo Fisher Scientific Cat# P36930).

### Explant Culture

From P0 *Phox2b^cre^;tdTomato* pups, both the left and right nodose ganglion were collected in Hibernate-A media (BrainBits, cat# HA) containing B27 supplement (ThermoFisher, cat# 17504044) and glutamax (ThermoFisher, cat# 35050061) and plated on poly-lysine coated chamber slides in a droplet of Growth Factor Reduced Matrigel (Corning, cat# 354230). Explants were incubated with a serum-free media containing Neurobasal media, B27 supplement, glutamax, and penicillin/streptomycin. To each well, either vehicle (PBS) or candidate factors were added at the following concentrations: BDNF (5ng/mL, R&D Systems, Cat# 248-BDB-005), GDNF (5ng/mL, ThermoFisher, Cat# RP-8602), NGF (50ng/mL, R&D Systems, Cat# 256-GF-100), LIF (10ng/mL, Gibco, Cat# PHC9484), and netrin (125ng/mL, R&D Systems, Cat# 6419-N1-025). Concentrations used are based on previously published work demonstrating effects of neurite outgrowth in the nodose ganglion and in other neural tissues (Schaller et al.; Niwa et al., 2002). Explant cultures were grown for 4 days in a standard 37°C incubator then imaged for analysis of neurite outgrowth.

### In situ hybridization

#### RNAscope

Paraformaldehyde fixed frozen samples from E13 and E15 whole mouse embryos were collected and sections containing the nodose ganglion and gastrointestinal tissue were processed for multiplex fluorescent *in situ* hybridization (RNAscope, ACD/Biotechne) using standard protocols specified by the manufacturer. Briefly, slides were baked for 1h at 60°C, post-fixed in 4% paraformaldehyde, and boiled in antigen retrieval reagent prior to hybridization with probes directed against mouse *Bdnf* or *Gdnf* mRNAs. Bound probes were detected using the RNAscope v2 multiplex fluorescent detection kit and Opal 520 and 690 Dyes (Akoya Biosciences).

#### Dual HiPlex RNAscope and Immunohistochemistry

Paraformaldehyde fixed frozen samples containing the right nodose ganglion were collected on E13, E15, P0, and P7 for processing for highly multiplexed mRNA labeling using the HiPlex RNAscope v2 platform from ACD. As differences have been reported between the left and right nodose ganglion, all histological analyses were performed solely on the right nodose ganglion for consistency (Han et al., 2018). Using a cryostat, 20um thick tissue sections were cut and mounted onto Superfrost Plus slides, baked at 60°C, and dehydrated in ethanol. Following antigen retrieval and protease treatments, sections were hybridized for 2 hours against the following 12 mouse probes: T1-*Scn10a*, T2-*Cckar*, T3-*Htr3b*, T4-*Ntrk2*, T5-*Gfra1*, T6-*Vip*, T7-*Calca*, T8-*Gpr65*, T9-*Oxtr*, T10-*Slc17a6*, T11-*Glp1r*, T12-*Sst*. Positive and negative control probe sets were run in parallel. Probe detection was performed in four rounds to detect T1-3, T4-6, T7-9, and T10-12 sequentially, accomplished with a confocal imaging step and fluorophore cleavage step between each round. A DAPI counterstain was also included in each round to serve as an anchor for later image registration.

Following detection of all probes, tissue was processed for immunohistochemistry to detect the boundaries of neuronal cell bodies. Slides were incubated for 30 minutes in blocking buffer containing 10% normal donkey serum (Jackson ImmunoResearch, West Grove, PA) and 1% bovine serum albumin (BSA) in PBS, then overnight in antibody solution containing 1% BSA in PBS with primary mouse anti-HuC/D antibody (ThermoFisher Scientific, Cat# A-21271, RID:AB_221448). Sections were washed two times for five minutes in PBS containing 0.1% Tween-20 (PBST), then incubated in secondary Alexa Fluor^®^ 594 AffiniPure F(ab’)2 Fragment Donkey Anti-Mouse IgG (Jackson ImmunoResearch Cat# 715-586-151, RRID:AB_2340858) antibody solution. After several rinses in PBST, sections were counterstained with DAPI (Thermo Fisher Scientific Cat# D1306), and a coverslip was applied using ProLong Gold Antifade Mountant (Thermo Fisher Scientific Cat# P36930). A final round of confocal imaging was then performed to capture HuC/D and DAPI labeling.

#### Image Acquisition

Leica STELLARIS or Zeiss LSM 710 laser scanning confocal microscopes equipped with 10x, 20x, 40x water-corrected, and 63x oil-corrected objectives were used to acquire fluorescence images. Confocal image stacks were collected through the z-axis at a frequency optimally determined by the imaging software based on the optics of the microscope and the wavelength of the fluorophores used for analysis. For all analyses, slides were coded so that the operator was blind to experimental group.

For analysis of activated Caspase+ and tdTomato+ neurons in the right nodose ganglion, every fifth consecutive section was imaged through the entire length of the ganglion using a 20x objective. For analysis of RNAscope labeling, the right nodose ganglion and intestinal tissue were identified and imaged using a 40x water-corrected objective. For analysis of HiPlex RNAscope labeling, two sections through the largest portion of the ganglion were captured in confocal image stacks using a 40x water-corrected objective. All images were collected on the Leica STELLARIS confocal microscope with the exception of the nodose ganglion explants which required the use of a longer working distance lens only available in our Cellular Imaging Core on the Zeiss LSM 710 confocal microscope. For explant images, the entire explant was imaged using a 10x objective and a z-step of 1um.

### Image Analysis

For cell counts of activated Caspase+ and tdTomato+ neurons in the nodose ganglion, all image stacks for each animal were manually analyzed using the ‘Cell Counter’ plugin within the FIJI/ImageJ software. The boundaries of the nodose ganglion and cell inclusion/exclusion criteria were agreed upon prior to cell counts. The narrowing of tissue between the jugular and nodose ganglion served as a morphological boundary for nodose ganglion cell counts. Only activated caspase colocalized with tdTomato was included in cell counts. This accounted for the vast majority of the caspase signal in the tissue (86%). Average profile diameter in each sample was measured and used to calculate estimated total cell counts in accordance with Abercrombie’s formula (Abercrombie, 1946).

Neurite outgrowth from nodose ganglion explants was quantified using FIJI/ImageJ software. Image stacks were projected through the z-axis, and signal was manually thresholded and binarized. Four neurite length measurements were made on each image in each cardinal direction from the edge of the explant to the tip of the neurite, for which values were averaged.

#### Single-Cell Automated Multiplex Pipeline for RNA Quantification and Spatial Mapping (SCAMPR Pipeline)

Following dual HiPlex RNAscope, immunohistochemistry, and imaging procedures as detailed above, confocal image stacks were analyzed using the SCAMPR pipeline (Ghoddousi et al., 2022). The SCAMPR pipeline facilitates analysis of highly multiplexed mRNA labeling at single neuron resolution. Briefly, image stacks corresponding to each tissue section were projected across the z-axis and aligned using ACD’s image registration software, with DAPI being the common signal used to anchor all images together. Automated cell segmentation was performed using the Cellpose algorithm in Google Colaboratory based on HuC/D labeling to identify neuronal soma boundaries (Stringer et al., 2021). ROI boundaries outlining each neuronal cell body were imported into FIJI/ImageJ and manually corrected for any inappropriately assigned cell profiles. Individual images for each mRNA probe signal were then processed in ImageJ to subtract background (rolling ball radius = 1) and thresholded to facilitate maximal detection of signal and removal of noise. Signal was binarized and area fraction of mRNA signal was measured for individual neuronal ROIs. Area fraction data was exported to a gene by cell matrix for further statistical analysis using GraphPad Prism and SCAMPR R scripts.

### Quantification and Statistical Analysis

Data were statistically analyzed and graphed using R or GraphPad Prism software (RRID:SCR_002798) and expressed as mean values ± standard error of the mean. Each mouse is considered a sample (or each ganglion in the case of explant cultures), with sample sizes for each group and statistical analysis used included in each figure legend. A D’Agostino-Pearson normality test was used to determine whether parametric or nonparametric statistical analyses should be performed. For data following a normal distribution, an ordinary one-way ANOVA was used to compare means. For data that failed to pass the D’Agostino-Pearson normality test, a nonparametric Kruskal-Wallis test (correcting for multiple comparisons using Dunn’s test) was used to compare mean rank difference. A p<0.05 was used for significance.

## Results

### Dynamic changes in gene expression in the developing nodose ganglion

To gain initial insight into how vagal sensory neurons are changing between early postnatal life and adulthood, RNA sequencing was performed comparing nodose ganglion samples from P7 and 12-week-old male mice. This revealed 1,453 genes were significantly increased and 1,734 genes were significantly decreased in adults (Figure 1A), representing more than 20% of the genes detected globally (3,188/14840). These data suggest that dramatic transcript-level changes in vagal sensory neurons take place following early postnatal life. By comparison, when adult male and female samples were analyzed using the same cutoffs, only 11 genes were found to be differentially expressed, suggesting little age-matched sexual dimorphism exists in the transcriptome of the mature nodose ganglion (data not shown).

**Figure 1.**
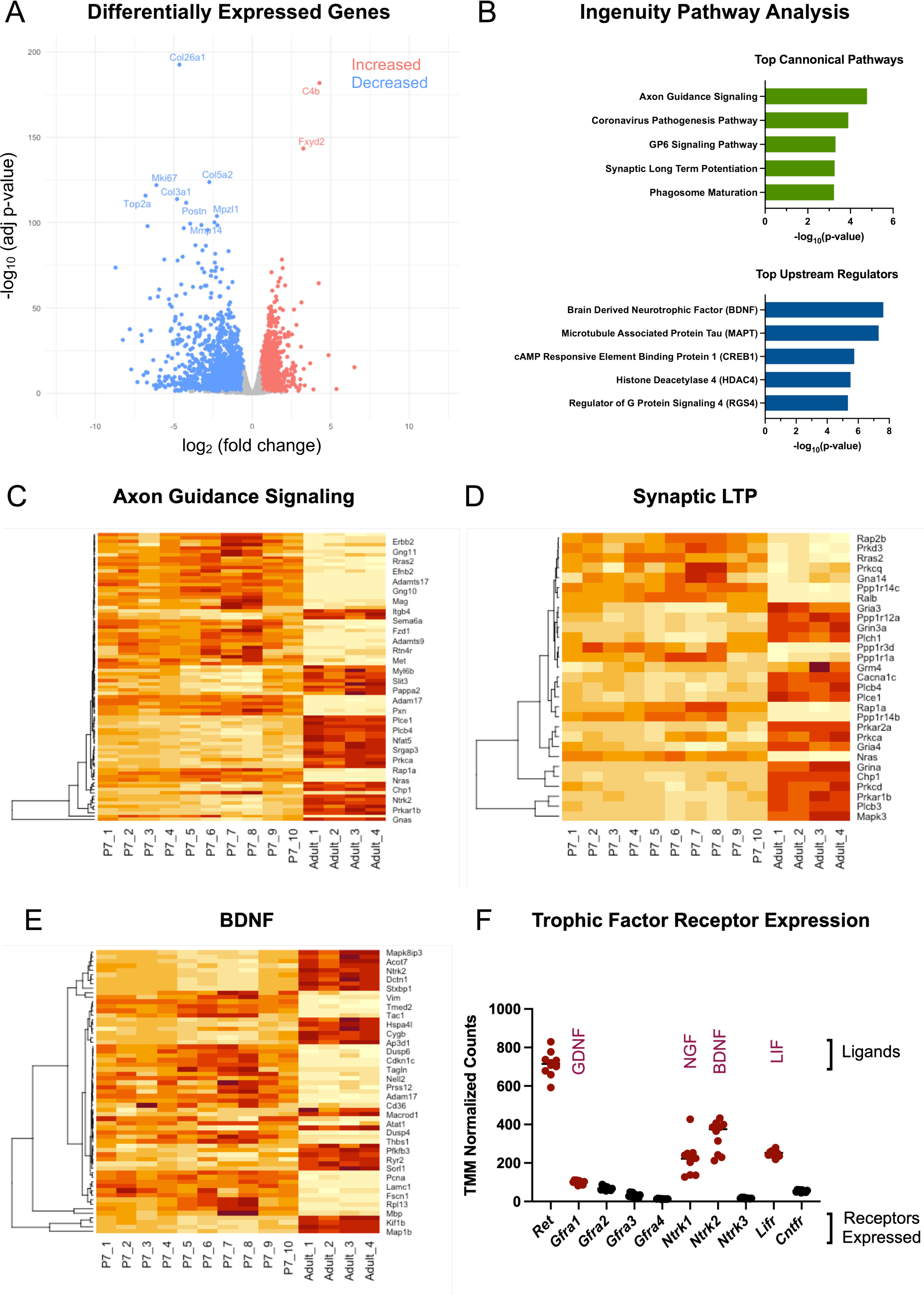
Transcriptomics analysis of nodose ganglion development reveals thousands of differentially expressed genes. (A) Volcano plot showing the 1,453 genes significantly increased (red, adjusted p-values <0.05, fold changes > 1.5) and 1,734 genes significantly decreased (blue, adjusted p-values <0.05, fold changes < −1.5) in adult male nodose ganglion samples compared to P7. (n = 10 P7 samples and 4 adult samples) (B) Ingenuity pathway analysis of DEGs revealed the top canonical pathways and top upstream regulators in early postnatal vs. adult samples. (C) Heatmap of DEGs related to axon guidance signaling. Warmer colors indicate higher gene expression. Individual samples are represented across columns. Specific genes are organized across rows grouped using hierarchical clustering. (D) Heatmap of DEGs related to synaptic LTP. (E) Heatmap of DEGs regulated by BDNF. (F) TMM normalized counts of trophic factor receptors expressed in P7 nodose ganglion samples. Values highlighted in red were the top 5 highest expressed on P7 and chosen for further testing in *in vitro* explant experiments.

**Figure 2.**
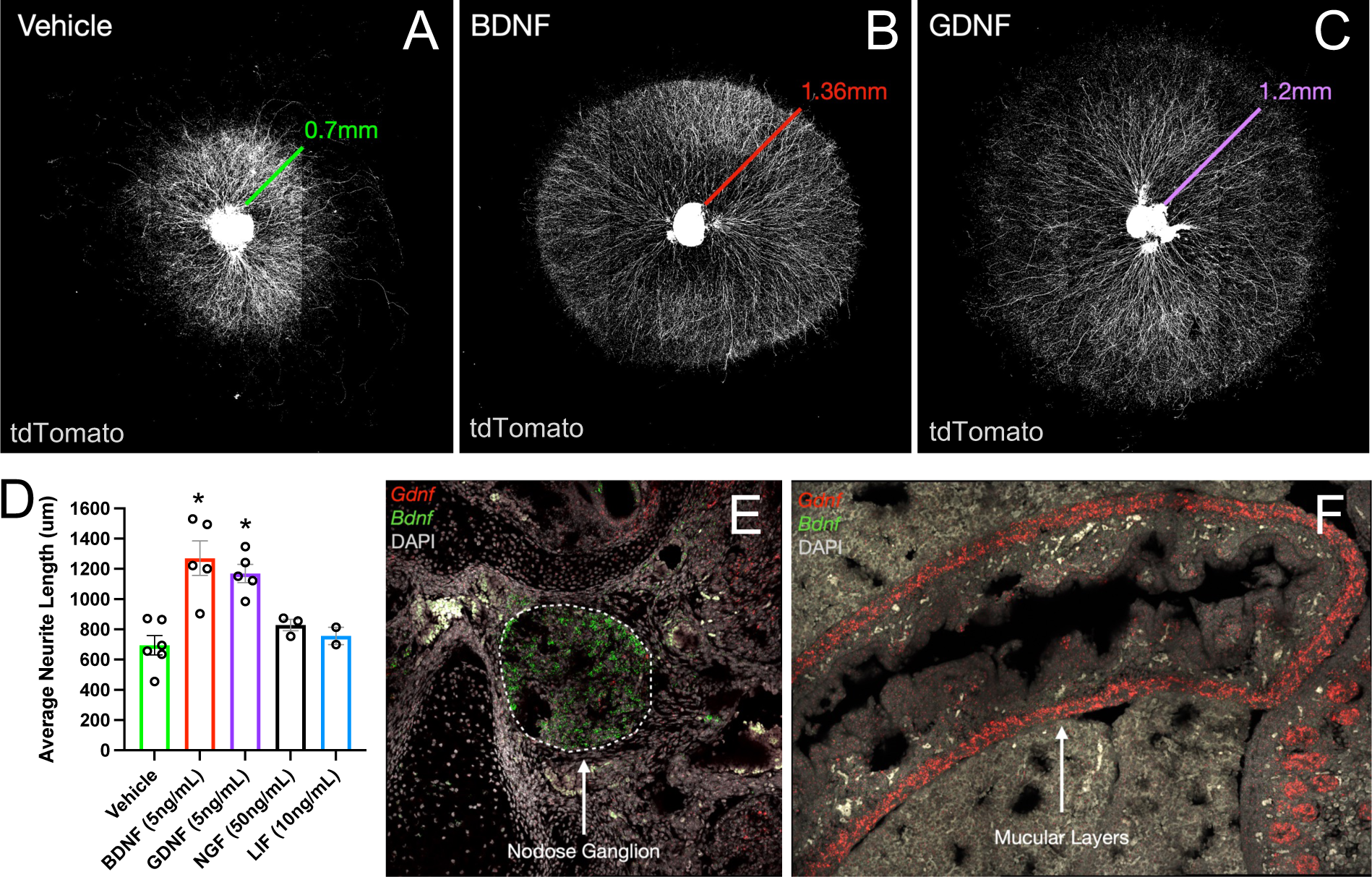
BDNF and GDNF stimulate neurite outgrowth from cultured nodose ganglion explants. (A-C) Representative images of nodose ganglion explants grown from *Phox2b^cre^;tdTom* mouse pups on P0 treated with vehicle (A), 5ng/mL BDNF (B), or 5ng/mL GDNF (C). Colored lines indicate average neurite lengths for each condition as graphed in (D). (D) Quantification of average neurite length of nodose ganglion explants grown for 4 days with vehicle, BDNF, GDNF, NGF, or LIF. * indicates p < 0.05 as analyzed by Kruskal-Wallis non-parametric ANOVA with Dunn’s correction for multiple comparisons. n = 6 vehicle, 5 BDNF, 5 GDNF, 3 NGF, 2 LIF. (E, F) Representative images of multiplex *in situ* hybridization for *Bdnf* (green) and *Gdnf* (red) and DAPI counterstain (grey) in E15 mouse embryos. Dashed lines in (E) outline the boundaries of the right nodose ganglion. Arrow in (F) points to *Gdnf* transcript labeling in the muscular layers of the intestine.

To further elucidate the biological implications of the considerable number of gene expression changes observed, QIAGEN Ingenuity Pathway Analysis was performed on the set of developmentally regulated differentially expressed genes (DEGs). Amongst the top canonical pathways identified, *Axon guidance signaling* and *Synaptic long-term potentiation* (Figure 1B-D) were amongst the top canonical pathways identified, consistent with potential continued circuit growth and synaptic strengthening between P7 and adulthood. Another pathway identified, the *coronavirus pathogenesis pathway*, was heavily composed of ribosomal protein genes known to be important for viral replication, but in this context may relate more closely to their critical function in the increased translation that is required for neurite outgrowth. Similarly, the DEGs enriched within the *GP6 signaling pathway* were primarily composed of collagen and laminin related genes that form the extracellular substrate that supports neurite outgrowth (refer to supplementary materials for full DEG and pathway lists). Among the top upstream regulators identified were *brain derived neurotrophic factor* (BDNF), which has known roles in neuronal growth and survival, and *microtubule associated protein tau* (MAPT), which functions to stabilize axonal microtubules (Figure 1B and E). Together, the sequencing data and pathway analysis demonstrate the substantial representation of genes involved in neurite growth, synaptic strengthening, and trophic factor signaling through BDNF.

BDNF is known to support the survival and growth of a subset of neurons in the nodose ganglion (Ernfors et al., 1994; Niwa et al., 2002; Hellard et al., 2004). This is in line with our transcriptomics data suggesting that BDNF is a significant regulator of developmental gene expression on P7. Further analysis of the sequencing data confirmed high expression of the BDNF receptor, *Ntrk2*, on P7 along with several other neurotrophic factor receptors (NTFRs) with well-established effects on nervous system development. Among the other most highly expressed NTFRs were *Ret* and *Gfra1*, which together bind glial derived neurotrophic factor (GDNF), *Ntrk1*, the receptor for nerve growth factor (NGF), and *Lifr*, the receptor for leukemia inhibitory factor (LIF; Figure 1F). Given the prominent representation of DEGs involved in axon growth and guidance in early postnatal life, we sought to examine the relationship between the highest expressed trophic factor receptors in early postnatal samples and growth of vagal sensory neurons.

### BDNF and GDNF support neurite outgrowth from the nodose ganglion

Using an in vitro system of nodose ganglion explants, candidate neurotrophic factors were tested for their ability to stimulate neurite outgrowth. To focus specifically on neurite outgrowth and reduce effects of neurotrophic factors on neuronal survival, ganglia were collected on P0 after the peak in apoptosis (Supplemental Figure 1). Ganglia cultured with vehicle survived well and extended neurites growing radially in all directions from the explant (Figure 1A). Compared to the vehicle treated explants, ganglia cultured with either BDNF or GDNF exhibited robust increases in average neurite length (Figure 1A-D). By contrast, ganglia cultured with NGF or LIF showed no difference in neurite length compared to vehicle treated cultures (Figure 1D). While differences in response properties and innervation patterns have been described in the left versus right nodose ganglion (Han et al., 2018), no differences in trophic factor responsiveness between sides was observed.

In addition to the explant culture system, a small number of dispersed cultures were tested in which nodose ganglion neurons were dissociated into single cells and plated onto laminin and poly-lysine coated wells. Unlike explant cultures where neuronal survival was favorable under both vehicle and neurotrophic factor treated conditions, vehicle treatment in dissociated cultures resulted in large scale neuronal death leaving few surviving neurons to test for trophic influences on neurite outgrowth. In dissociated cultures, BDNF treatment significantly improved neuronal survival. However, as the vehicle condition could not be used to examine neurite outgrowth, the dissociated culture model was abandoned in favor of explant cultures. The difference in survival between vehicle treated explants and vehicle treated dispersed cultures suggests that local factors produced within the nodose ganglion are important for sustaining neuronal survival.

As BDNF and GDNF were both found to increase neurite outgrowth from developing nodose explants, their expression patterns were determined during the perinatal period using multiplex fluorescent *in situ* hybridization (RNAscope). Interestingly, BDNF was highly expressed in the E15 nodose ganglion itself (Figure 1E) in a pattern that overlapped with the neuronal marker, HuC/D. Conversely, little GDNF was expressed in the nodose ganglion. Instead, GDNF was highly expressed in the muscular layers of the GI tract, and modestly expressed in stromal and epithelial compartments of the intestine (Figure 1F). Together, the data show that while both BDNF and GDNF support nodose neurite outgrowth, BDNF may act locally within the nodose ganglion itself and GDNF may support target derived growth of ganglion projections to subdiaphragmatic targets in the GI tract.

### GDNF receptor expression is enriched in subdiaphragmatic projecting Scn10a+ neurons

Responsiveness to BDNF and GDNF are dependent upon expression of their respective neurotrophic factor receptors; *Ntrk2* for BDNF and *Gfra1* for GDNF. While binding of GDNF to *Gfra1* also requires expression of *Ret*, *Ret* dimerizes with other receptors such as *Gfra2*, *Gfra3*, and *Gfra4* to confer specificity of binding to other trophic factors (neurturin, artemin, and persephin, respectively) and is therefore not specific to GDNF (Airaksinen and Saarma, 2002). The enrichment of GDNF observed in the GI tract raises the question of whether expression of the GDNF receptor, *Gfra1*, is also enriched in vagal sensory neurons that project to the stomach and intestine. To this end, highly multiplexed *in situ* hybridization (HiPlex RNAscope) was used to label 12 independent RNA transcripts within the same tissue section. This facilitated labeling of several cell-type specific nodose ganglion markers with *Ntrk2* or *Gfra1*. Tissue samples were collected from mice on E13, E15, P0 and P7 to examine trophic factor receptor expression as nodose sensory neurons grow and reach GI targets, and as they mature in postnatal life. Analysis across these ages demonstrated that *Ntrk2* is expressed in the majority of vagal sensory neurons at all pre- and postnatal ages examined (Figure 3 AD). By contrast, *Gfra1* was expressed by a significantly higher fraction of nodose ganglion neurons at E13 than E15 or P7. Despite this decline in the proportion of neurons expressing *Gfra1*, *Gfra1* was expressed at higher levels in the cells that continued to express it (Figure 3A-D).

**Figure 3.**
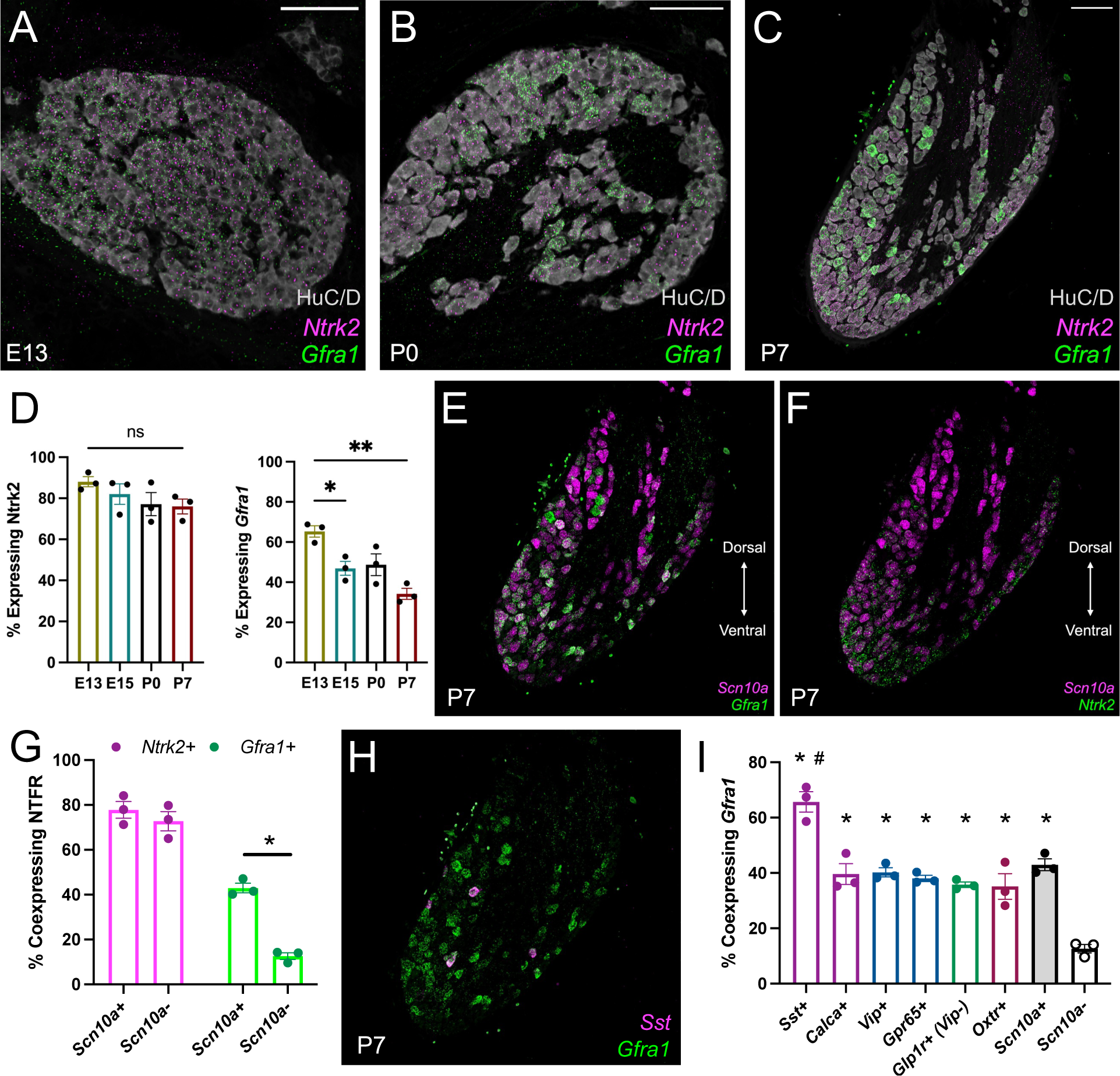
GDNF receptor expression is enriched in *Scn10a*+ vagal afferent cell types that project to the GI tract. (A-C) Representative images of the right nodose ganglia on E13 (A), P0 (B), and P7 (C) showing labelled *Gfra1* (green) and *Ntrk2* (magenta) transcripts, or the neuronal marker HuC/D (grey). (D) Quantification of the proportion of neurons expressing *Ntrk2* or *Gfra1* on E13, P0, and P7. (n = 3 mice per group). * indicates p < 0.05, *** indicates p < 0.001, ‘ns’ indicates not significant as analyzed by one-way ANOVA and Tukey’s correction for multiple comparisons. (E, F) Representative images of labelled *Scn10a* (magenta), *Gfra1* (green, E) and *Ntrk2* (green, F) transcripts in the right nodose ganglion of P7 mouse pups. (G) Quantification of the percentage of *Scn10a*+ and *Scn10a*- that co-express *Ntrk2* or *Gfra1* on P7. (n = 3 mice per group). * indicates p < 0.05 as analyzed by 2-sided t-test. Statistics for *Ntrk2* and *Gfra1* were performed separately, then graphed together. (H) Representative image of labelled *Sst* (magenta) and *Gfra1* (green) transcripts in the right nodose ganglia of a P7 mouse pup. (I) Quantification of the percentage of neurons expressing each cell type marker (x-axis) that coexpress *Gfra1* on P7. Values for *Scn10a*+ and *Scn10a*- neurons also appear in (G) and are illustrated on the same graph for direct comparison. (n = 3 mice per group). * indicates p < 0.05 compared to *Scn10a*- percent coexpression by one-way ANOVA and Tukey’s correction for multiple comparisons. # indicates p < 0.05 compared to *Scn10a*+ percent coexpression by one-way ANOVA and Tukey’s correction for multiple comparisons.

**Figure 4.**
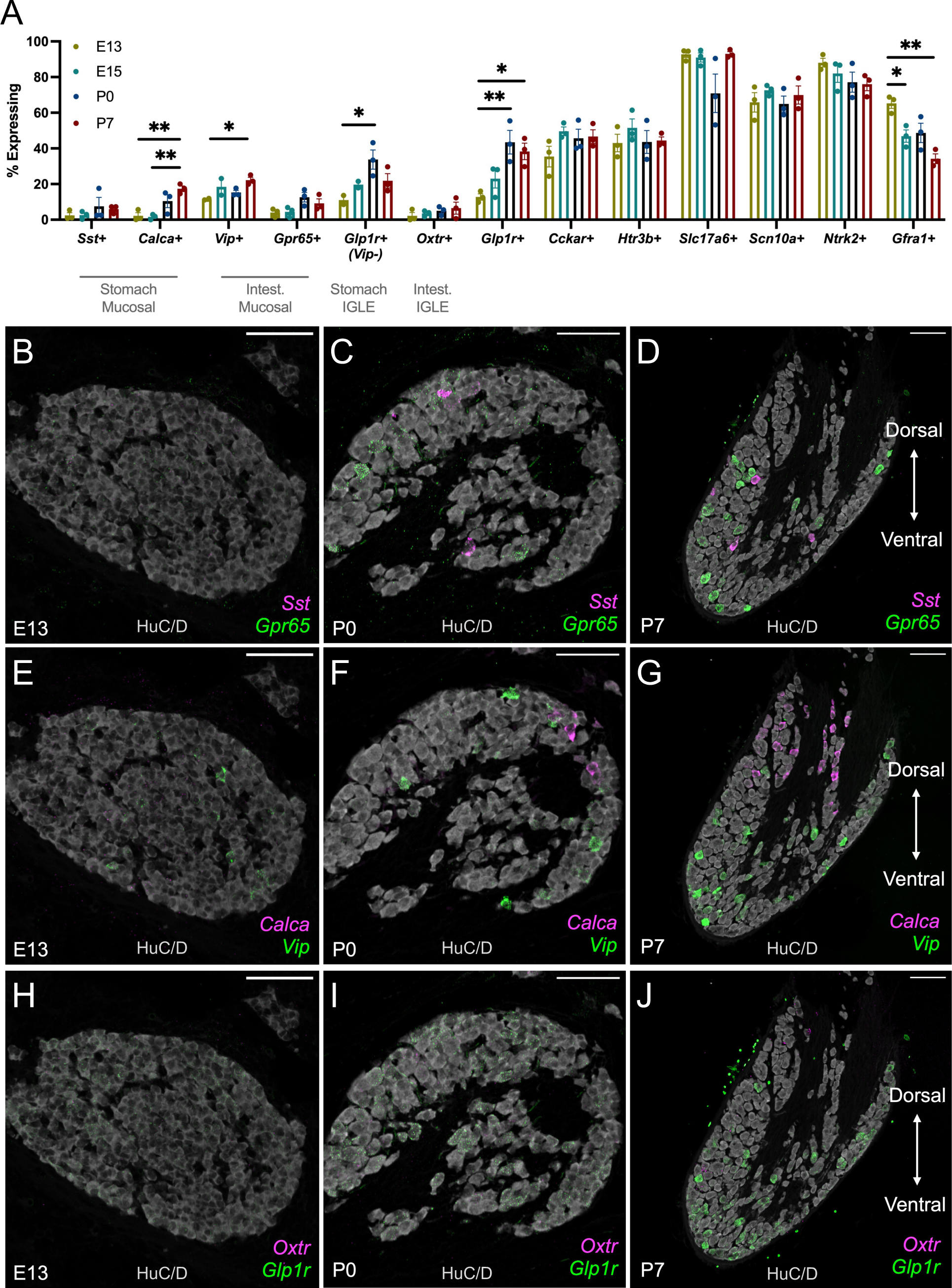
Early prenatal expression of vagal afferent cell type markers becomes progressively more mature by P7. (A) Quantification of all 12 transcripts measured using HiPlex RNAscope and the SCAMPR pipeline for analysis at single neuron resolution. mRNAs probed are indicated on the x-axis. The percentage of neurons expressing each transcript is indicated on the y-axis. Each point represents percentages calculated from a single mouse, in which expression was quantified from several hundred neurons in the right nodose ganglion. (n = 3 mice per group; 324 - 795 neurons measured per sample). * indicates p < 0.05 comparing all ages for each transcript by one-way ANOVA and Tukey’s correction for multiple comparisons. (B-D) Representative images of the right nodose ganglion on E13, P0, and P7 labelled for *Sst* (magenta) and *Gpr65* (green) transcripts and the neuronal marker, HuC/D. Scalebar = 100um. (E-G) Representative images of the right nodose ganglion on E13, P0, and P7 labelled for *Calca* (magenta) and *Vip* (green) transcripts and HuC/D. Scalebar = 100um. (H-J) Representative images of the right nodose ganglion on E13, P0, and P7 labelled for *Oxtr* (magenta) and *Glp1r* (green) transcripts and HuC/D. Scalebar = 100um.

To identify the subpopulation of neurons expressing *Gfra1* or *Ntrk2*, each NTFR was examined for colocalization with specific vagal cell type markers on P7. Anatomical and single cell sequencing studies have demonstrated that *Scn10a* (also known as Nav1.8) is primarily expressed by vagal sensory neurons that project below the diaphragm (~90%), with a minor portion of afferents projecting to pulmonary targets (Bai et al., 2019; Kupari et al., 2019b). On P7, *Ntrk2* was expressed by a similar proportion (approximately 75%) of *Scn10a*+ and *Scn10a*- neurons on P7 (Figure 3F-G). By contrast, only 12% of *Scn10a*- neurons expressed *Gfra1*, while 43% of Scn10a+ neurons co-expressed *Gfra1*, demonstrating a significant enrichment of *Gfra1* in a neuronal population that projects heavily to the GI tract (Gautron et al., 2011; Bai et al., 2019; Kupari et al., 2019b) (Figure 3E, G). A distinct enrichment of *Gfra1*+ neurons was apparent in the ventral portion of the fused nodose-jugular ganglion, corresponding to expression in the ventral nodosal compartment. These expression patterns are consistent with a role for target derived GDNF expressed by the developing GI tract in supporting subdiaphragmatic vagal growth.

Subdiaphragmatic *Scna10a*+ vagal cell types can further be subtyped by expression of other marker genes. Stomach mucosal projecting neurons express either *Sst* or *Calca*, stomach intraganglionic laminar endings (IGLEs) express *Glp1r* but not *Vip*, intestinal mucosal projecting neurons express either *Gpr65* or *Vip*, and intestinal IGLEs express *Oxtr* (Figure 5B; (Bai et al., 2019)). Using HiPlex RNAscope, each of these cell types were examined for co-expression of *Gfra1* on P7. Compared to *Scn10a*- neurons, all GI projecting populations (*Sst*+, *Calca*+, *Vip*+, *Gpr65*+, *Glp1r*+/*Vip*-, and *Oxtr*+) exhibited a significant enrichment of *Gfra1* expression (Figure 3I), with *Sst*+ stomach mucosal neurons exhibiting the highest enrichment of *Gfra1* compared to all *Scn10a*+ neurons (Figure 3H-I).

**Figure 5.**
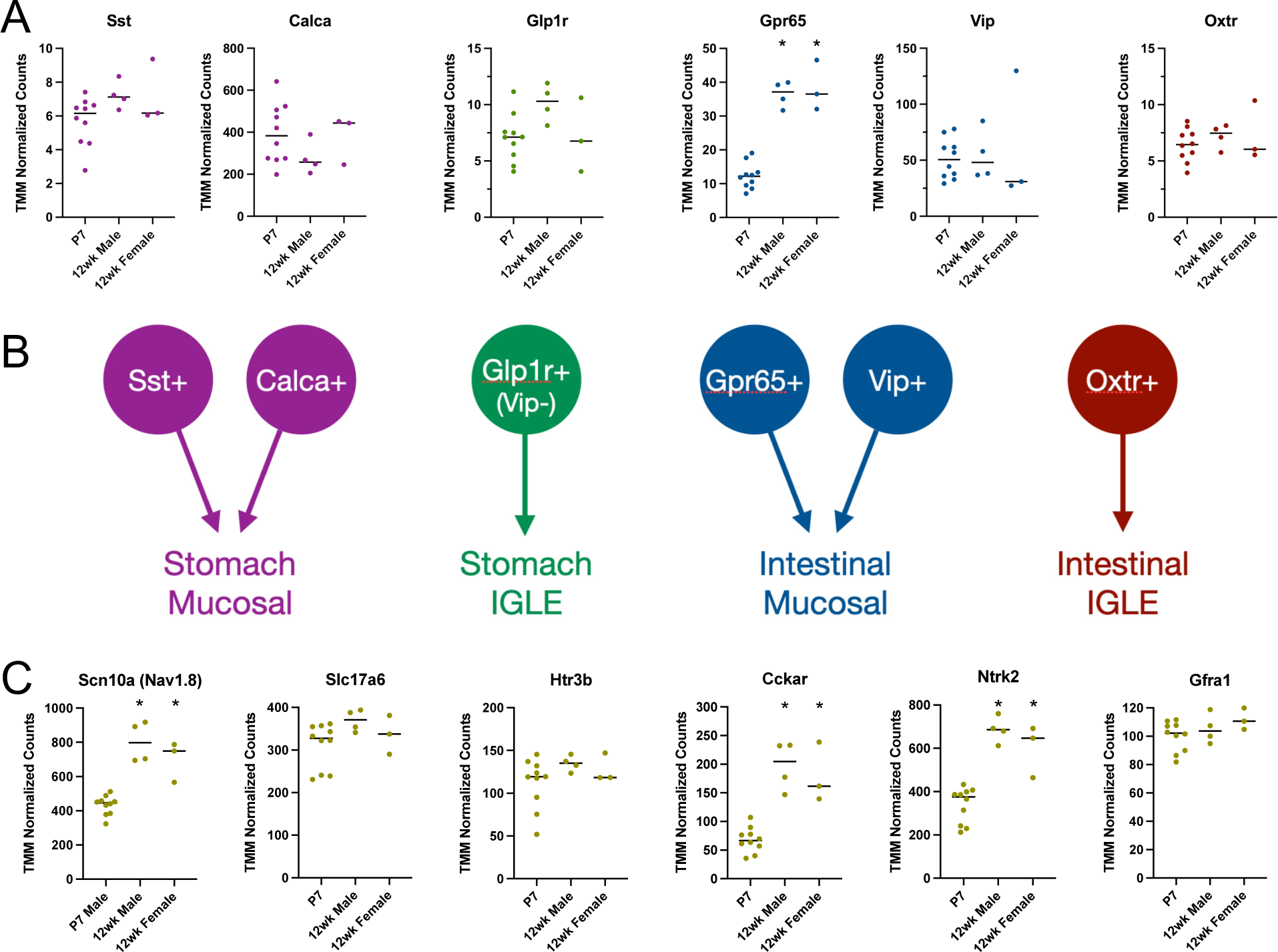
Expression of some vagal afferent cell type markers continues to increase after the first postnatal week. (A) TMM normalized counts of vagal afferent cell type markers expressed in P7 male and adult male and female nodose ganglion samples. (n = 10 P7 male samples, 4 adult male samples, 3 adult female samples). (B) Schematic depicting each cell type marker and its respective subtype definition as projecting to the stomach vs. intestine, and as an IGLE vs. mucosal afferent neuron type. (C) TMM normalized counts of other notable cell defining markers expressed in P7 male and adult male and female nodose ganglion samples. Markers graphed are the same ones probed in HiPlex RNAscope experiments described in Figure 4. (n = 10 P7 male samples, 4 adult male samples, 3 adult female samples). * indicates p < 0.05 comparing all groups by one-way ANOVA and Tukey’s correction for multiple comparisons.

Potential *Ntrk2* and *Gfra1* enrichment in select subpopulations of nodose ganglion neurons was also analyzed in embryonic samples. At E13, a similar proportion of *Scn10a*+ and *Scn10a*- neurons coexpressed *Gfra1* (approximately 65%, Supplementary Figure 2). By E15, coexpression of *Gfra1* was significantly decreased in *Scn10a*- neurons (52.8% of *Scn10a*+, 31.1% of *Scn10a*-). This difference was sustained in P0 (57.9% of *Scn10a*+, 31.4% of *Scn10a*-) and P7 (43.0% of *Scn10a*+, 12.6% of *Scn10a*-) nodose ganglion. A significant, but transient, decrease in the proportion of *Scn10a*- neurons expressing *Ntrk2* was also observed on E15, but was not sustained in P0 or P7 nodose ganglion. While analysis of coexpression within *Scn10a*+ neurons was possible at embryonic time points, coexpression of NTFR in specific subpopulations of GI-projecting neurons was not possible due to low expression of most cell type marker genes. Instead, expression patterns were leveraged to examine maturation of cell type marker expression.

### Nodose cell types are readily resolved by early postnatal life

Nodose ganglion cell type identity and function has primarily been examined in adult rodents (Gautron et al., 2011; Egerod et al., 2018; Bai et al., 2019; Kupari et al., 2019b). As such, there are limited data available that describe the development of vagal sensory transcriptomic diversity and cell type identity. To establish a timeline for vagal afferent cell type specific marker expression, HiPlex RNAscope and the SCAMPR analysis pipeline were used. On E13 and E15, little to no signal was concentrated within specific cell bodies for *Sst*, *Calca*, *Gpr65*, or *Oxtr* transcripts (Figure 4A, B, E, H; representative E15 ganglia images not shown, but quantified in 4A). Conversely, dense *Vip* signal was observed in a small number of cell bodies at E13 (Figure 4E-G). The proportion of *Vip*+ neurons increased through E15 and P7 (Figure 4A, E-G). Additionally, neurons expressing the serotonin receptor, *Htr3b*, also exhibited a dense expression pattern in a subpopulation of neurons that was maintained by a constant proportion of neurons from E13 to P7 (Figure 4A, Supplementary Figure 3A-C). The expression patterns of *Vip* and *Htr3b* in specific neuronal subpopulations on E13 suggests that, while still immature, the process of defining specific cell types has begun by this time point. By P7, concentrated labeling for all cell type markers were observed localized to specific single neuron profiles (Figure 4D, G, J, Supplementary Figure 3C).

A dorsal-ventral specific labeling pattern was observed for several transcripts on P7. *Sst*, *Gpr65*, *Vip*, and *Glp1r* were primarily localized to the ventral portion of the nodose-jugular ganglion complex (Figure 4D, G, and J). Conversely, *Calca*+ neurons were localized to the dorsal portion of the ganglion (Figure 4G). While *Calca* is expressed by ~10% of gastric projecting nodose neurons, it is also expressed by *Prdm12*+ neurons of the jugular ganglion, which are located in the dorsal most portions of the fused jugular-nodose ganglion (Bai et al., 2019; Kupari et al., 2019b).

Examination of RNA sequencing data was further used to compare changes in cell type marker gene expression between P7 male and adult male and female. No statistically significant differences in cell type marker expression were observed by comparing adult male and female nodose ganglion. Compared to adult ganglia, there was no change in expression for *Sst*, *Calca*, *Glp1r*, *Vip*, or *Oxtr*, suggesting that by P7 a mature expression pattern has been reached for these genes in nodose neurons (Figure 5A). By contrast, *Gpr65*, a marker of a subtype of intestinal mucosal neurons, increased more than 3-fold in adult male and female nodose ganglion, suggesting that this population continues to mature after P7 (Figure 5A). GPR65 is a membrane bound G protein-coupled receptor able to detect changes in extracellular pH and is present in macronutrient responsive populations of vagal neurons (Williams et al., 2016a). *Scn10a*, *Cckar*, and *Ntrk2* were also increased in adults (Figure 5C). The voltage-gated sodium channel, Scn10a (Nav1.8), modulates vagal electrophysiological properties, while the CCK receptor, Cckar, participates in vagally mediated satiety signaling and meal termination (Gibbs et al., 1973; Raybould and Tache, 1988; Moran, 2000). The expression data for *Gpr65*, *Scn10a*, and *Cckar* suggest that the response properties of GI projecting vagal neurons continue to mature in early postnatal life as they are exposed to feeding related stimuli.

## Discussion

Cell type definitions established in adults through both genomic and functional studies have been instrumental to understanding vagal sensory communication and physiological regulation of ingestive behavior. The present study leverages these cell type definitions to further understand the timing of when vagal cell types emerge developmentally and their responsiveness to trophic influences. Specifically, we demonstrate that both BDNF and GDNF stimulate neurite outgrowth in vagal sensory neurons. Mapping of neurotrophic factor expression patterns developmentally indicates that while BDNF may act locally to support growth from the nodose ganglion itself, GDNF may act as a target derived growth factor within the muscular layers of the intestine. Consistent with the latter, the GDNF receptor, *Gfra1*, is enriched preferentially in *Scn10a*+ neurons that primarily project below the diaphragm to GI sites, and in all of the vagal cell types known to project to the GI tract. Lastly, highly multiplexed *in situ* hybridization data demonstrate that discrete cell type marker expression in specific subpopulations of nodose neurons can be detected as early as E13 for transcripts such as *Vip* and *Htr3b*, suggestive of specification of cell types by this age. Patterns of cell type marker expression, however, are largely immature prenatally, and further develop into discretely labelled subpopulations by the end of the first postnatal week.

### Transcriptomic data indicate large differences in gene expression in early postnatal life compared to adults

Transcriptomics analyses revealed that an astonishing 20% of genes in the nodose ganglion are differentially expressed between P7 and adults. Heavily represented among these DEGs were transcripts involved in axon growth and guidance. This was somewhat surprising given past descriptions suggesting that on P7 vagal sensory circuit morphology in the GI tract is similar to patterns observed in adults (Murphy and Fox, 2007). Similarly, central projections of vagal axons to terminal regions in the nucleus of the solitary tract have been described as mature as early as P2 in rats (Rinaman and Levitt, 1993). Despite the relatively mature pattern of innervation observed in early postnatal life morphologically, smaller scale increases in neurite outgrowth and in refinement at synapses may be at play. This is further supported by the enrichment of DEGs associated with synaptic LTP. It is clear from seminal behavioral studies that the ability for nutrient related post-ingestive feedback to modulate feeding does not arise until after the second postnatal week (Phifer et al., 1986; Weller, 2000, 2006; Weller et al., 2000; Smith, 2006). Therefore, it is possible that some of the expression changes identified in the present study may relate to ongoing maturation and synaptogenesis in circuits required for nutrient detection.

### Neurotrophic factor and receptor expression patterns

*In situ* hybridization experiments demonstrate that while BDNF is expressed by neurons of the nodose ganglion themselves, GDNF is expressed by distal targets in the GI tract. Together these data suggest that BDNF acts as a locally derived signal at the level of the nodose ganglion itself, while GDNF works instead as a target derived signal to support vagal neuron growth and survival. This hypothesis is additionally supported by in vitro experiments demonstrating that vagal neuronal survival is low in dissociated cultures, and high in explant cultures where the cells are packed tightly together, suggesting that a factor produced within the ganglia itself improves neuronal survival. While additional experiments are necessary to confirm this, it is likely that keeping the ganglia intact facilitates the local action of BDNF on neuronal survival. Exogenous BDNF enhances explant outgrowth to a greater extent than vehicle alone, clearly demonstrating its importance in vagal sensory neurite growth. The exact subcellular site where BDNF is released remains to be determined as release from the cell body versus from centrally or peripherally projecting processes could have discrete functional consequences.

At all developmental ages tested, the BDNF receptor (*Ntrk2*) was expressed by the vast majority of neurons (78-88%), suggesting its broad importance for most vagal neurons. By comparison, the pattern of expression of *Gfra1* was such that while it was expressed more widely but at lower levels in prenatal life, it became more highly expressed in a smaller subset of only 34% of neurons by P7. *Gfra1* was additionally shown to be enriched in *Scn10a*+ populations compared to *Scn10a*- populations, and was similarly enriched amongst several functionally defined cell types, particularly in *Sst*+ neurons. Single cell sequencing studies performed in adults suggest that *Gfra1* is prevalent in *Gpr65*+ populations (Kupari et al., 2019b). Our results indicate no greater enrichment in *Gpr65*+ neurons compared to any other GI projecting population, however, it is noteworthy that the sequencing data in the present study indicate a more than 3-fold increase in *Gpr65* expression that occurs between P7 and adulthood. It is possible that that this enrichment of *Gfra1* in *Gpr65*+ neurons occurs later in development after the first postnatal week.

### Timing of cell type marker expression

Initial expression of specific marker genes is a clear indication that cell type has been specified. Our multiplex *in situ* hybridization data demonstrate that as early as E13, transcripts for *Vip* and *Htr3b* are already highly expressed within specific subpopulations of neurons. The expression patterns of marker genes continue to mature considerably on through postnatal life, however, it is remarkable how early this process begins. In adult rodents, *Vip*+ neurons project exclusively to the mucosal compartment of the small intestine (Bai et al., 2019). Developmental studies demonstrate that vagal sensory axons first reach the intestine between E12 and 14, and begin to innervate the mucosal compartment between E14 and 16 (Ratcliffe et al., 2006; Hao et al., 2020; Niu et al., 2020). The timing of these developmental events together with the marker expression data from the current study demonstrate that defined *Vip*+ subpopulations arise either prior to, or concurrent with reaching their terminal innervation targets.

In contrast to the data for *Vip*, most of the other marker genes did not exhibit restricted cell type specific expression patterns until later time points. It is an important distinction to note that the appearance of the cell type markers used in the present study is only one measure of neuronal maturation. Some genetic markers may be expressed either earlier or later than other markers in neurons that are at comparable developmental states. For example, cell type may already be specified by E13 in stomach mucosal neurons despite not expressing high levels of *Sst* or *Calca*. Further experiments will be required to more precisely define when these cell fate decisions occur for all vagal subpopulations. However, the *Vip* and *Htr3b* data along with the caspase3 results in the current study suggest that some cell type specification has occurred already by E13 even as other developmental processes such as continued neurite outgrowth and apoptosis proceed. Other published studies indicate that by P0, there are morphologically distinct projections within the GI tract characteristic of IGLEs, IMAs, and mucosal endings, suggesting that some vagal sensory cell types are clearly defined by birth (Murphy and Fox, 2007). Our data confirm and extend this timeline, suggesting that defined cell type emerge prenatally.

In addition to the early onset of cell-type specific expression observed for some marker genes, prolonged maturation in expression was also noted for other marker genes in the bulk RNA sequencing dataset. Notably *Cckar* and *Gpr65* significantly increased in expression between P7 and 12-weeks. Both of these genes are important for nutrient detecting vagal pathways. CCKA receptors detect CCK released from intestinal enteroendocrine cells in response to nutrient ingestion and *Gpr65*+ neurons respond electrophysiologically to nutrient ingestion (Williams et al., 2016b; Kaelberer et al., 2018). The finding that their expression increases more than 3-fold after P7 is consistent with other functional studies that have demonstrated that the ability of ingested nutrients to modulate feeding behavior develops after the second postnatal week. The continued postnatal maturation of functional properties is interesting and raises the question of whether early life nutritional environment could modulate these developmental trajectories. Future work will be required to further parse these important developmental events.

## Acknowledgements

The image acquisition work was performed in the Cellular Imaging Core at The Saban Research Institute (TSRI), Children’s Hospital Los Angeles (CHLA). The RNA Sequencing work was performed in the Single Cell, Sequencing, and CyTOF (SC2) Core at TSRI, CHLA. Experiments were supported by a Stem Cell Challenge Award from the Keck School of Medicine, University of Southern California and funds from the Developmental Neuroscience and Neurogenetics Program at CHLA. We thank students Andrea Lanz and Elizabeth Huber from the Samuels Family Latino and African American High School Internship Program for their work quantifying explant data.

---

**Supplemental Figure 1.**
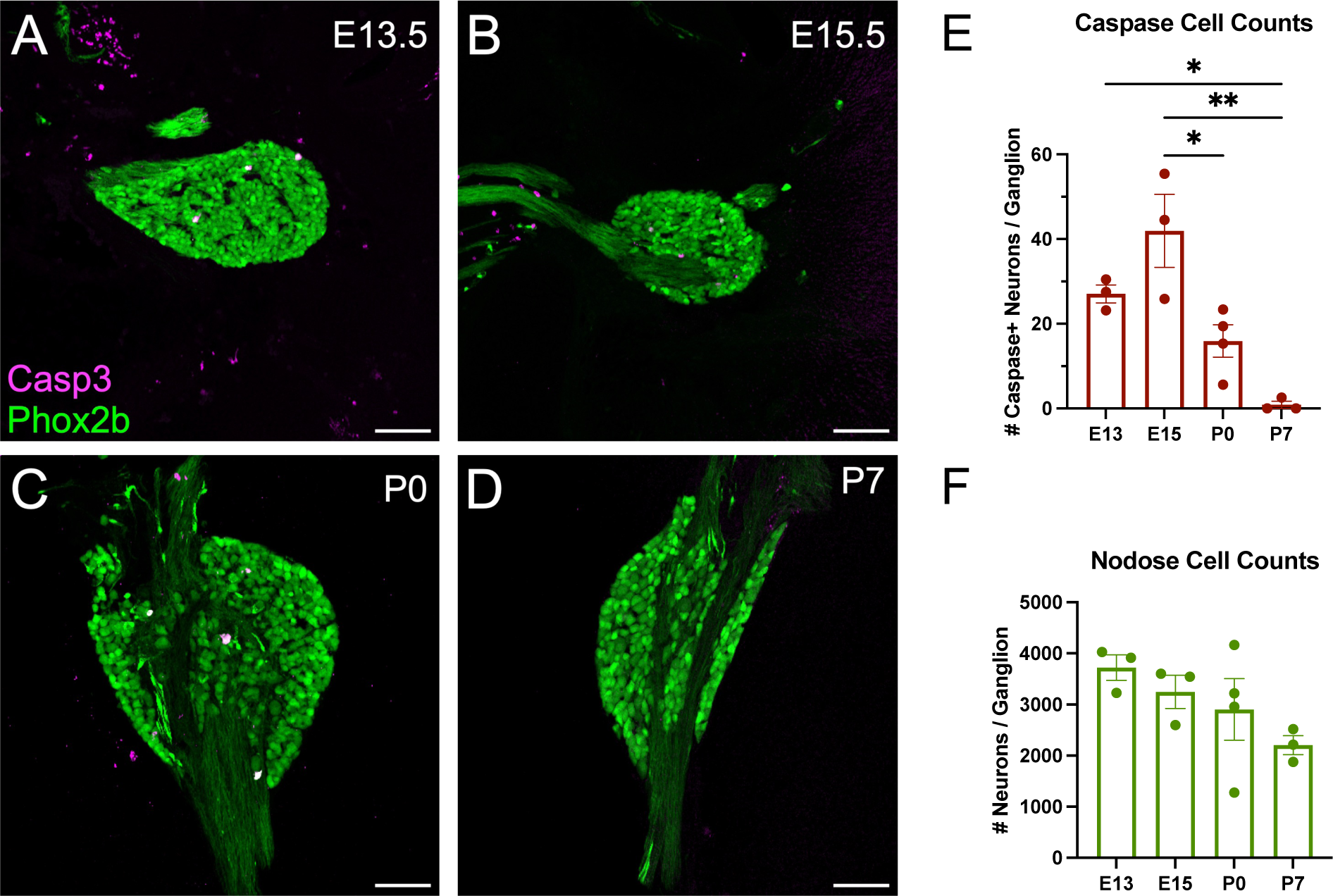
Ontogeny of cell death in the nodose ganglion. (A-D) Representative images of the right nodose ganglion on E13, E15, P0, and P7 from *Phox2b^cre^;Tomato* mice. Immunohistochemistry was used to label the apoptotic marker, activated caspase 3 (magenta), together with endogenous tdTomato labeling (green) of all vagal sensory neurons. (E) Quantification of the number of activated caspase positive neurons per ganglion on E13, E15, P0 and P7. (n = 3 to 4 mice per group). * indicates p < 0.05, ** indicates p < 0.01 comparing all groups by one-way ANOVA and Tukey’s correction for multiple comparisons.

**Supplemental Figure 2.**
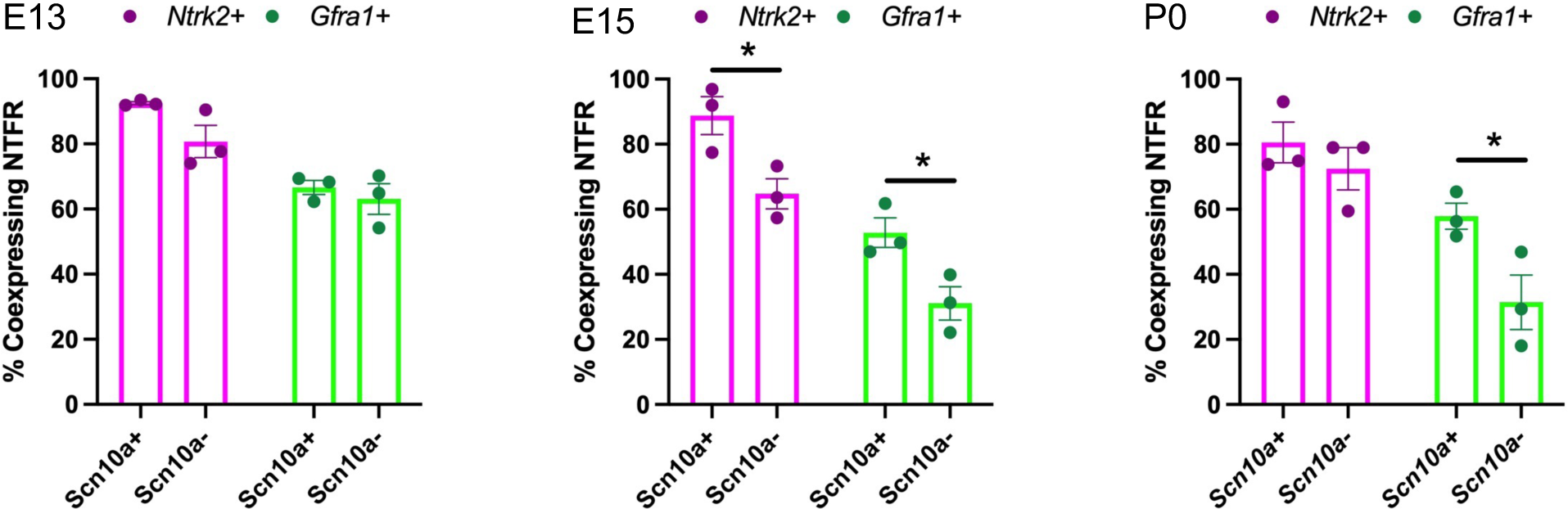
Neurotrophic factor receptor expression in *Scn10a*+ and *Scn10a*- neurons across development. The percentage of neurons coexpressing either *Ntrk2* or *Gfra1* was measured on E13, E15, and P0. (n = 3 mice per group). * indicates p < 0.05 as analyzed by 2-sided t-test at each age. Statistics for *Ntrk2* and *Gfra1* were performed separately, then graphed together.

**Supplemental Figure 3.**
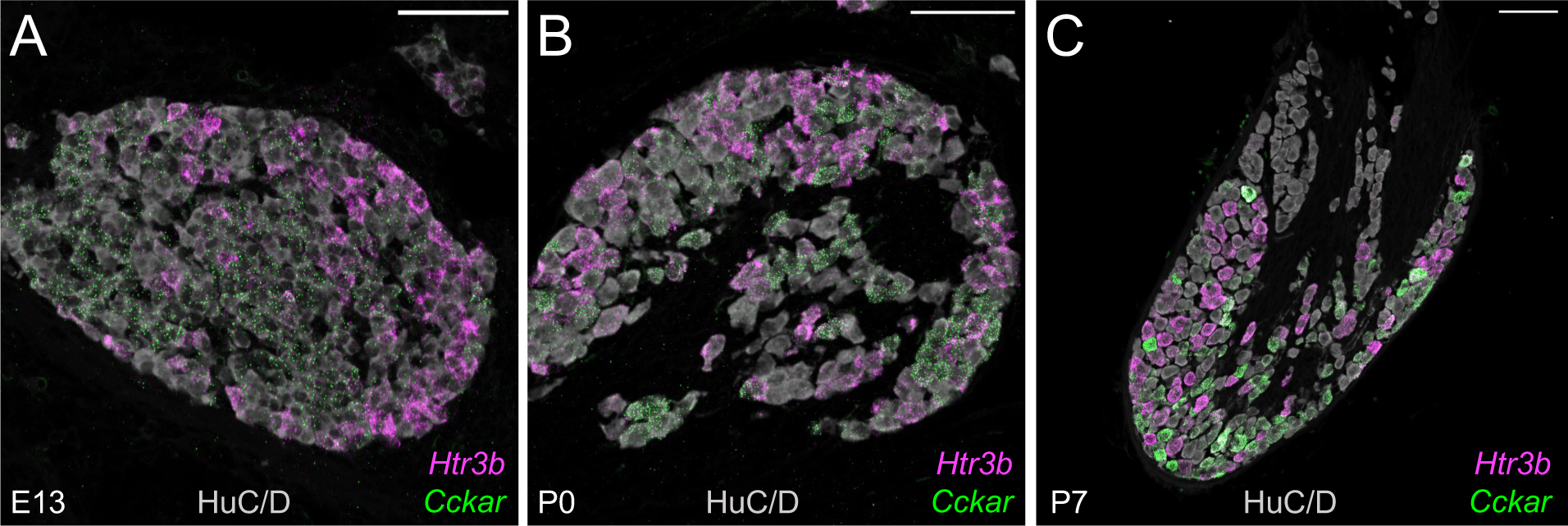
Early specification of subpopulations of vagal neurons expressing *Htr3b*. (A-C) Representative images of the right nodose ganglion on E13, P0, and P7 labelled for *Htr3b* (magenta) and *Cckar* (green) transcripts and HuC/D. Scalebar = 100um.

